# HIV-1 specifically traps CD9 and CD81 tetraspanins within viral buds and induces their membrane depletion

**DOI:** 10.1101/293860

**Authors:** Selma Dahmane, Christine Doucet, Antoine Le Gall, Célia Chamontin, Patrice Dosset, Florent Murcy, Laurent Fernandez, Desirée Salas Pastene, Eric Rubinstein, Marylène Mougel, Marcelo Nollmann, Pierre-Emmanuel Milhiet

**Author notes:** These two authors equally contributed to the work (Selma Dahmane, Christine Doucet).

## Abstract

HIV-1 assembly specifically alters both partitioning and dynamics of the tetraspanins CD9 and CD81 forming enriched areas where the virus buds. Importantly the presence of these proteins at exit sites and in viral particles inhibits virus-induced membrane fusion. To get molecular insights into tetraspanins partitioning in this viral context, we correlated nanoscale CD9 mapping obtained by super resolution microscopy to membrane topography probed by Atomic Force Microscopy (AFM). We demonstrated that CD9 is specifically trapped within the nascent viral particles, especially at buds tips, and that Gag mediate CD9 and CD81 depletion from cellular surfaces, even in the absence of Vpu and Nef, resulting from tetraspanins escaping from the plasma membrane during HIV-1 release. In addition, we showed that CD9 is organized as small membrane assemblies of few tens of nanometers that can coalesce upon Gag expression. Our results support a functional redundancy among tetraspanins during HIV release.

## AUTHOR SUMMARY

As an obligate enveloped virus, Human immunodeficiency type 1 virus (HIV-1) subverts the plasma membrane to enter and escape from the host cell. Apart from viral proteins, the release of HIV-1 requires specific cellular components, which are recruited to the egress site by the viral protein Gag. In this study, we used a combination of advanced microscopies to investigate how Gag affects host membrane proteins, the tetraspanins CD9 and CD81 that are key players in many infectious processes including HIV-1 replication. It is currently unclear how tetraspanins influence Gag assembly and how Gag assembly alters tetraspanin organization at the plasma membrane. Here we correlated super resolution microscopy with membrane topography delineated by the sharp tip of an atomic force microscope (AFM) on cells expressing HIV-1 Gag. We report for the first time high-resolution mapping of tetraspanin organization during virus egress and found that Gag proteins drastically reorganize and recruit these proteins within both nascent and mature budding membrane structures. We also observed that Gag expression promote tetraspanin depletion at the cell surface, which is likely due to tetraspanin escape from the plasma membrane when virions are released.

## INTRODUCTION

Human Immunodeficiency type 1 virus (HIV-1) egress from host cells is a key process in its dissemination in infected patients, which is initiated by the structural polyprotein Gag that is necessary and sufficient to release virus-like particles (VLPs)(1). Gag is expressed as a 55 kDa polyprotein composed of 4 structural domains (matrix (MA), capsid (CA), nucleocapsid (NC) and p6) that will be cleaved after particle release. After its synthesis in the cytosol, Gag is targeted to the inner leaflet of the plasma membrane where it multimerizes, inducing the formation of membrane curvature (budding sites) and finally membrane fission by recruiting host factors such as ESCRT machinery (2). The N-terminally myristoylated MA domain targets the polyprotein to the plasma membrane and lipids of the host plasma membrane have early been proposed to play a key role in this process (3), especially acidic lipids such as phosphatidylserine (PS) and phosphatidylinositol biphosphate (PI(4,5)P2), as well as sphingolipids and cholesterol (4,5). The latter two lipids are known to form or be enriched in different types of microdomains, especially raft microdomains that could behave as pre-formed recruitment platforms (4). Because raft microdomains had been described as ordered membrane domains (6,7), it was proposed that HIV-1 Gag proteins can sense cholesterol and acyl chain environment in membranes. In contrast, it was recently shown that HIV assembly does not simply result from a higher affinity of Gag to ordered membrane domains, but could involve an additional protein machinery (8). Proteins of the tetraspanin family could be part of it.

Tetraspanins belong to a family of proteins characterized by four transmembrane regions and a specific fold in the larger of the 2 extracellular domains. All human cell types express several of these proteins which play an essential role in multiple cellular processes ranging from cell morphology, migration, cell-cell fusion and signalling (9). Tetraspanins have been proposed to be molecular organizers and/or scaffolding proteins within the plasma membrane forming a dynamic network of protein-protein interactions at the cell surface by interacting with one another and with other transmembrane proteins (integrins, Immunoglobulin superfamily proteins and others) (10-12), referred to as the tetraspanin web or Tetraspanin-enriched microdomains (TEM)(13). A fraction of tetraspanins and associated proteins concentrate into microscopically visible structures named tetraspanin-enriched areas (TEA) or platforms (14,15).

Tetraspanins are involved in several infectious diseases including Plasmodium falciparum, Listeria monocytogenes and viruses such as hepatitis C virus (HCV) and HIV-1 (16,17). Notably, previous studies have shown colocalization of several tetraspanins (CD9, CD63, CD82 and CD81) with HIV-1 Gag and Env, in several cell types including T cells (18-20). Critically, TEMs were proposed to constitute gateways for HIV-1 assembly and budding. More recently, we have demonstrated using single molecule tracking experiments that both CD9 and CD81 are specifically recruited and sequestered within Gag assembly sites, supporting the idea that cellular and viral components, instead of clustering at pre-existing microdomain platforms, direct the formation of distinct domains enriched in tetraspanins for the execution of specific functions, yet not fully elucidated (21).

Studies based on tetraspanin knockdown or inhibition by specific antibodies also revealed that tetraspanin down-regulation decreases virus entry and replication in macrophages (22,23) In addition several studies pinpointed a potential role of tetraspanins in modulating HIV-1 infectivity through their incorporation into the released viral particles. Overexpression of tetraspanins in virus-producing cells led to the production of virions with less infectivity (24,25). The presence of these proteins at exit sites also reduced the formation of syncitia in producing cells and cell-to-cell fusion induced by the virus (24,26,27). Conversely, CD81 and CD82 levels were shown to be downregulated by the HIV-1 accessory proteins Vpu and Nef, which induce protein sequestration in intra-cellular compartments, leading to decreased levels at the plasma membrane, and degradation (28). These observations raise important questions concerning the role of CD9 and CD81 in the different steps of viral replication, from the recruitment of Gag polyproteins at the plasma membrane to the budding and release of viral particles.

Here, we show that Gag mediates depletion of both CD9 and CD81 from the surface of cells. This happens even in the absence of the regulatory proteins Vpu and Nef and depends on VLPs release. Moreover, this phenomenon was not observed for non-related plasma-membrane proteins. Altogether, this suggests that CD9 and CD81 depletion is due to their accumulation in Gag-induced VLP budding sites. To understand how they accumulate, we characterized tetraspanins distribution by super resolution microscopy (29). We show that CD9 organizes as small nanoscale units and follows the distribution of Gag. Correlative dSTORM/AFM further shows that the tetraspanin concentrates within nascent viral particles. In most cases, it localizes at the very tip of viral buds and is excluded from the bases of the budding sites. Combination of these two nanoscopy techniques thus demonstrates that CD9 trapping by Gag is restricted to viral buds. This supports our proposed mechanism for tetraspanin depletion from host cells. Overall, our data reveal how Gag, by modulating tetraspanin distribution and expression, impacts the hot cell environment. Our results also suggest a redundancy among tetraspanins. This could explain the difficulties to assess the role of tetraspanins during HIV-1 life cycle.

## RESULTS

### Gag reduces the cell surface expression of both tetraspanins CD9 and CD81

Previous studies clearly established that TEMs serve as exit sites for HIV-1 and we have demonstrated that Gag assembly specifically induced the formation of tetraspanin-enriched platforms at the surface of HeLa cells (21). As observed in T lymphocytes (25), this recruitment could induce incorporation of tetraspanins within released virions and we then analyzed CD9 and CD81 expression in HeLa cells expressing or not Gag tagged with GFP (named Gag-GFP). Cells were transfected with equimolar ratios of pGag and pGAG-GFP, inducing the biosynthesis of VLPs mimicking HIV-1 infection (30), and tetraspanins were stained with labeled anti-CD9 or anti-CD81 antibodies. Confocal micrographs strongly suggest that CD9 and CD81 levels decreased in Gag-expressing cells as compared to control cells (Fig. 1A) and this trend was confirmed by measuring the cell surface expression of both tetraspanins by flow cytometry. 30-40% of cells were positively transfected and we defined 3 types of populations: untransfected cells (GFP-) and cells with intermediate (GFP+) or high (GFP++) levels of expression (Fig. 1B). While GFP-transfected cells had comparable CD9 levels in the 3 populations, CD9 surface levels were decreased by 70-80% in cells strongly expressing Gag- GFP, as compared to GFP-negative cells from the same sample. Intra-sample ratios of CD9 and CD81 levels in GFP++ or GFP+ versus GFP- cells were averaged from 4 independent experiments (Fig. 1C) and the values confirmed that cells with high Gag expression are depleted of CD9 and CD81. In contrast, we found that the cell surface levels of CD46, a non-raft transmembrane protein with little association with tetraspanins, remains unaffected upon overexpression of Gag proteins (data not shown), indicating that Gag specifically affects CD9 and CD81 surface levels.

**Figure 1.**
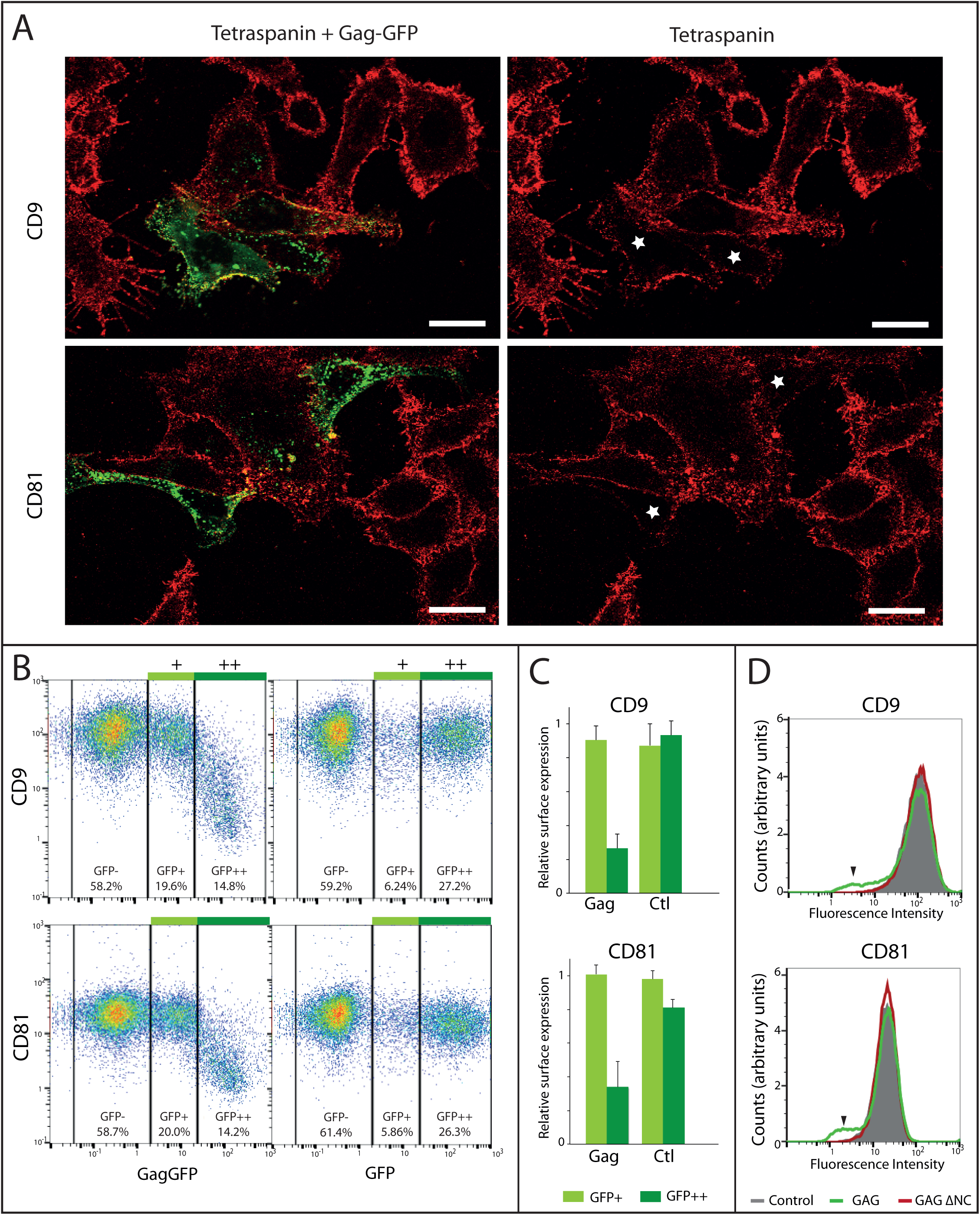
Gag reduces tetraspanin levels at the cell surface of HeLa cells. **A)** Confocal images of HeLa expressing Gag-GFP proteins. CD9 and CD81 are labeled with antibodies functionalized with Alex647. White stars indicate cells where tetraspanins are downregulated as compared to Gag-GFP negative cells. Scale bar is 25 μm. **B, C and D)** Surface expression of CD9 and CD81 measured by flow cytometry 48h after transfection with Gag-GFP or GFP (Control). B) Representative flow cytometry 2D dot plots of gated living cells; three cell populations were defined based on their GFP intensity (GFP-, GFP+ and GFP++). Percentages indicate the representative fraction of each population; C) Mean CD9 and CD81 levels were quantified for each population and normalized to GFP- levels. Data were averaged from 3 independent experiments. Error bars are standard deviations. D) Surface levels of CD9 and CD81 in HeLa cells transfected with GFP alone (grey), with Gag-GFP (green line) or with the Gag ΔNC mutant (red line). Black arrowheads indicate HeLa cells with CD9 or CD81 depletion.

We then hypothesized that this depletion was due to the high excision rate of membrane buds enriched in CD9 and CD81. To test this, we used a Gag mutant protein, where the nucleocapsid domain was deleted (GagΔNC). This mutant assembles at the plasma membrane but is impaired in VLPs release (31). When expressed in HeLa cells, GagΔNC did not reduce tetraspanin cell surface levels (Fig. 1D). Altogether, fluorescence microscopy and flow cytometry indicate that HIV-1 Gag overexpression promotes downregulation of the surface expression of tetraspanin proteins CD9 and CD81 and this is due to tetraspanin escape from the plasma membrane when VLPs are released.

### Nanoscale organization of CD9 during Gag assembly

In order to precisely determine how tetraspanins and Gag are orchestrated during the formation of HIV-1 viral particles, we first performed direct stochastic optical reconstruction microscopy (dSTORM) imaging using TIRF illumination to compare the membrane organization of CD9 in control and Gag-expressing HeLa cells (32). Cells were transfected with pGAG-GFP (see above) and then stained with anti-CD9 antibodies labelled with Alexa647. Importantly, all experiments were performed in fixed cells since tetraspanins have been shown to be very dynamic (10). The localization precision was below 30 nm (Fig. S1A). dSTORM images of non-transfected control HeLa cells displays a sparse distribution of CD9 molecules on the basal membrane surface (Fig. 2A, left column). The mean density of CD9 localizations *(i.e*. detected events) in control cells was 1192 μm^2^ ± 127 (sem) (Fig. 2B and Table S1). Upon Gag expression (Fig. 2A, second and third columns and Fig. S1B), Gag-GFP foci assembled at the plasma membrane (white), consistent with previous reports, and CD9 partitioned into Gag-enriched areas (Fig. 2A). The mean density of localizations in cells 24h and 48h after transfection with pGAG-GFP decreased to 793 ± 141 and 658 ± 151 per μm^2^, in agreement with the reduction in CD9 expression upon Gag expression described in Figure 1. Interestingly, using a mask based on Gag-GFP signal, we noted a dramatic increase in the localization density within Gag- GFP-enriched areas at the cost of surrounding areas (8430 ± 1655 versus 428 ± 63 localizations/μm^2^ in areas devoid of Gag, 48h after transfection) (Fig. 2B and Table S1). As expected from Fig. 2A, CD9 localization density in Gag-GFP foci was correlated to Gag-GFP intensity (Fig. S1C, Fig. S1D, and Table S4 for Kendall's tau correlation coefficients).

**Figure 2.**
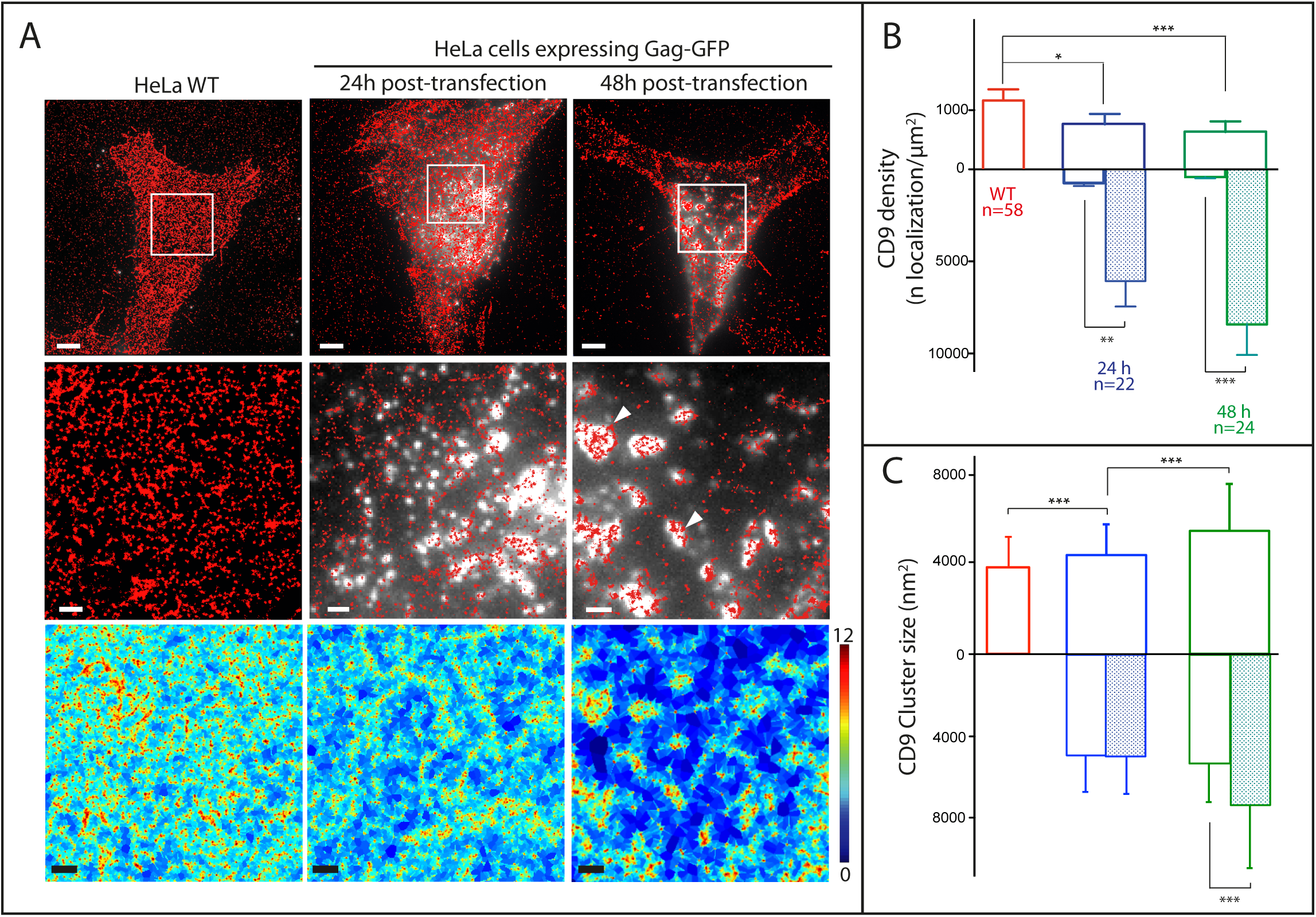
Gag proteins recruit and reorganize the tetraspanin CD9 at the plasma membrane. **(A)** Raw CD9 dSTORM localizations (red dots, 2 first rows) and local density map obtained from the SR-Tesseler framework (third row; the colour scale represents local densities in logarithmic scale) in HeLa cells expressing HIV-1 Gag-GFP (white signal in micrographs). From left to right: control cells; cells expressing Gag-GFP for 24h or 48h; the middle row show zoomed areas outlined in the upper images. Scale bars are 5 μm (upper row) and 1 μm (bottom rows); **(B)** dSTORM analysis. CD9 density (number of localization/μm^2^) in control cells (red) or in cells expressing Gag-GFP proteins 24h (blue) or 48h (green) after transfection. In the mirror histograms below the X axis, empty and hatched histograms represent the density outside and within Gag-GFP positive areas, respectively. Error bars are SEM and n is the number of analyzed cells. *, ** and *** indicate p values below 0.05, 0.001 and 0.0001 respectively, as determined by the Mann–Whitney U-test (for exact p values, see Table S1); **(C)** Histograms of size distribution of CD9 clusters in nm^2^ for the 3 conditions. The legend is similar to B.

To refine our analysis of CD9 lateral reorganization, single molecule localizations were analyzed using a segmentation procedure based on Voronoï diagrams, allowing a precise and automatic segmentation and quantification of protein organization (33)(Fig. 2A). This framework named "SR-Tesseler" was used to estimate the size of CD9 clusters at the plasma membrane expressed in nm^2^ (for more details, see the Supplemental Materials section). In control cells, the mean area of CD9 clusters or assemblies was 3710 ± 1513 nm^2^ (Fig. 2C and Table S2) that correspond to a disk of 68.7 nm diameter. A significant increase of this mean area was observed upon Gag expression, up to 5471 ± 2198 nm^2^ for cells analyzed 48h after transfection that corresponds to a disk of 83.5 nm diameter.

This increase was even more pronounced when considering only Gag-enriched areas using the mask method described above: 6527 ± 2763 nm^2^ for cells 48h post-transfection versus 3710 ± 1513 nm^2^ for control cells, corresponding to a disk of 91.2 diameter (see Table S2 and the mirror histograms in Fig. 2C). These results demonstrate that CD9 trapping in Gag-enriched areas led to an accumulation of protein clusters in the 70-90 nm diameter range in average that can eventually coalesce to form larger membrane platforms. As a consequence, large areas depleted of CD9 were observed (Fig. 2A, 48h panel, and the quantitation of depleted areas Fig. S1E).

Taken together we observe that Gag expression in host cells leads to a dramatic change in CD9 organization at the plasma membrane, forming larger clusters that concentrate into Gag assembly sites.

### Correlation between Gag assembly, CD9 recruitment and membrane curvature

To further understand the mechanism of Gag assembly and how CD9 recruitment and membrane budding are coordinated, we implemented correlative TIRF and AFM imaging. To ensure that apical cellular membranes imaged by AFM were in the TIRF evanescent field, we focused on thin regions present at the cell periphery where apical membranes imaged by AFM were in the TIRF evanescent field (Fig. 3). Topographic images of the cell surface revealed membrane protrusions that overlapped with Gag-GFP foci, suggesting they were VLPs. Some fluorescent areas did not coincide with membrane protrusion, likely because Gag assembly occurred on the basal membrane not accessible to the AFM tip (see the asterisk in Fig. 3). Virus-like buds were on average 104 ± 49 nm high and 162 ± 75 nm wide, which is in accordance with measurements derived from super resolution microscopy (see the review (34) or electron microscopy studies (35). As expected the size distribution of viruslike buds was heterogeneous (Fig. S2A), reflecting the progression of the budding process. As a matter of fact, the surface of buds measured by AFM correlates with their Gag-GFP content (Fig. 3E). Gag-GFP intensity could thus be used as a “molecular timer” to chronologically compare bud maturation and CD9 recruitment. In addition, AFM allowed to further characterize VLPs formation as shown by the section in Figure 3F where the AFM tip could delineate 2 budding sites that cannot be differentiated with conventional TIRF microscopy. This illustrates well the gain in resolution obtained by AFM.

**Figure 3.**
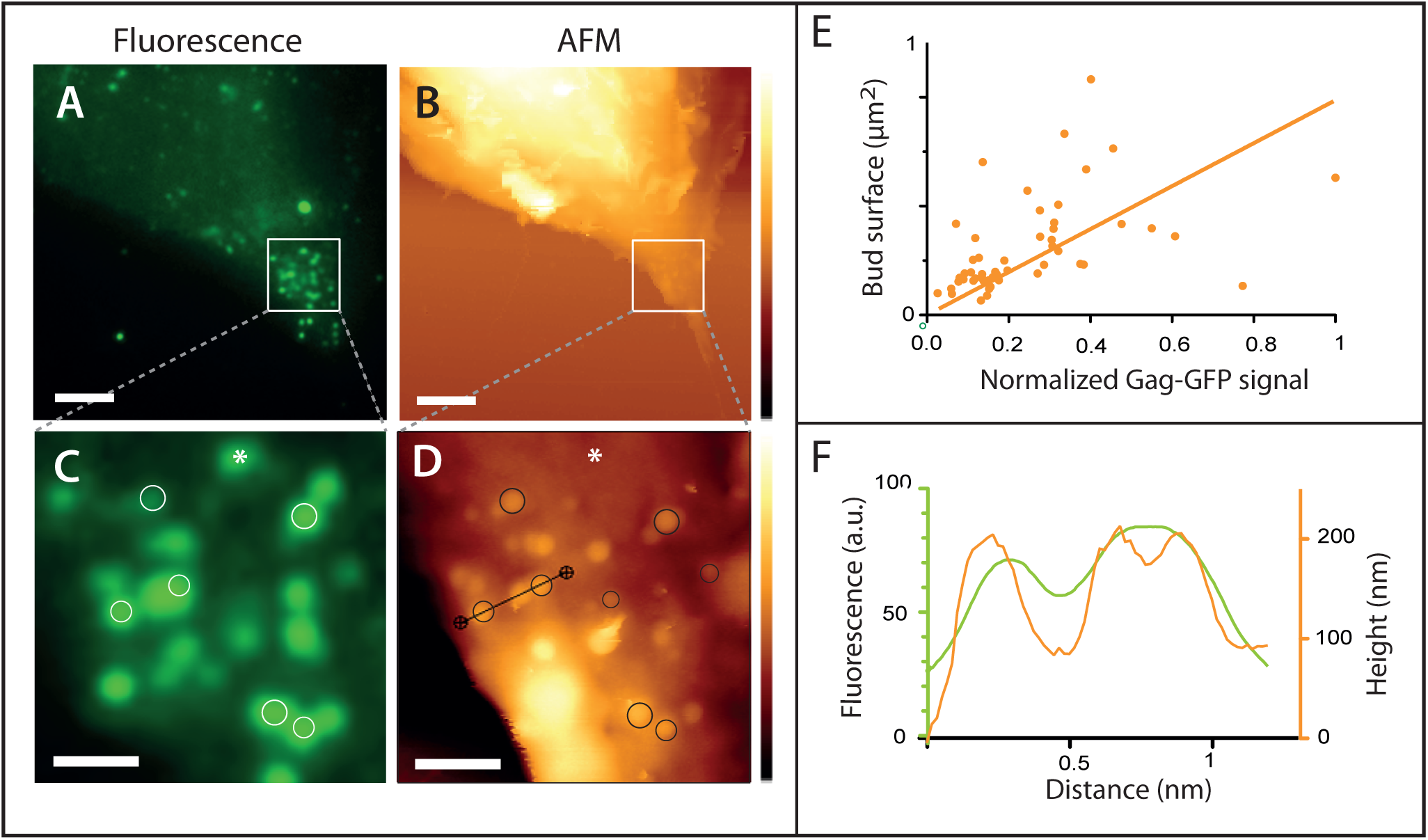
Nanoscale imaging of HeLa cells expressing Gag-GFP using correlated fluorescence-atomic force microscopy. Gag-GFP fluorescence image (A and C) and AFM topographic images (B and D) were compared (48h post-transfection here). The circles highlight some correlation between fluorescence and the membrane protrusion delineated by the AFM tip. (E) Normalized Gag-GFP signal as a function of the bud surface measured by AFM. (F) Profile plot of the topography (orange line) and fluorescence signal (green line) along the section indicated by the black line on the AFM image. The asterisk points out a Gag assembly where no membrane protrusion was observed by AFM. Scale bars are 5 (A and B) or 1 μm (C and D). The AFM z colour scales are 6.6 μm (upper AFM image) and 800 nm (lower image).

### CD9 dSTORM mapping within VLP buds

We thus combined dSTORM and AFM on cells expressing Gag-GFP to get more details on CD9 organization in Gag-enriched domains (Fig. 4). CD9 clusters characterized by dSTORM overlapped well with the shape of Gag-GFP budding sites delineated by the AFM tip (Fig. 4A to 4D). In advanced buds (spherical bud attached to the membrane by a neck), the neck area was devoid of CD9 (e.g. Fig. 4C). In nascent buds CD9 was localized at the very tip of membrane protrusions and mostly excluded from budding sites basis (Fig. 4F and see the gallery of budding sites in Fig. S3). This demonstrates that CD9 is indeed trapped within the nascent bud. Moreover, CD9 seems to preferentially associate with membrane regions of high positive curvature. As described in Fig. 2, some CD9 clusters were also observed in membrane areas devoid of Gag-GFP buds.

**Figure 4.**
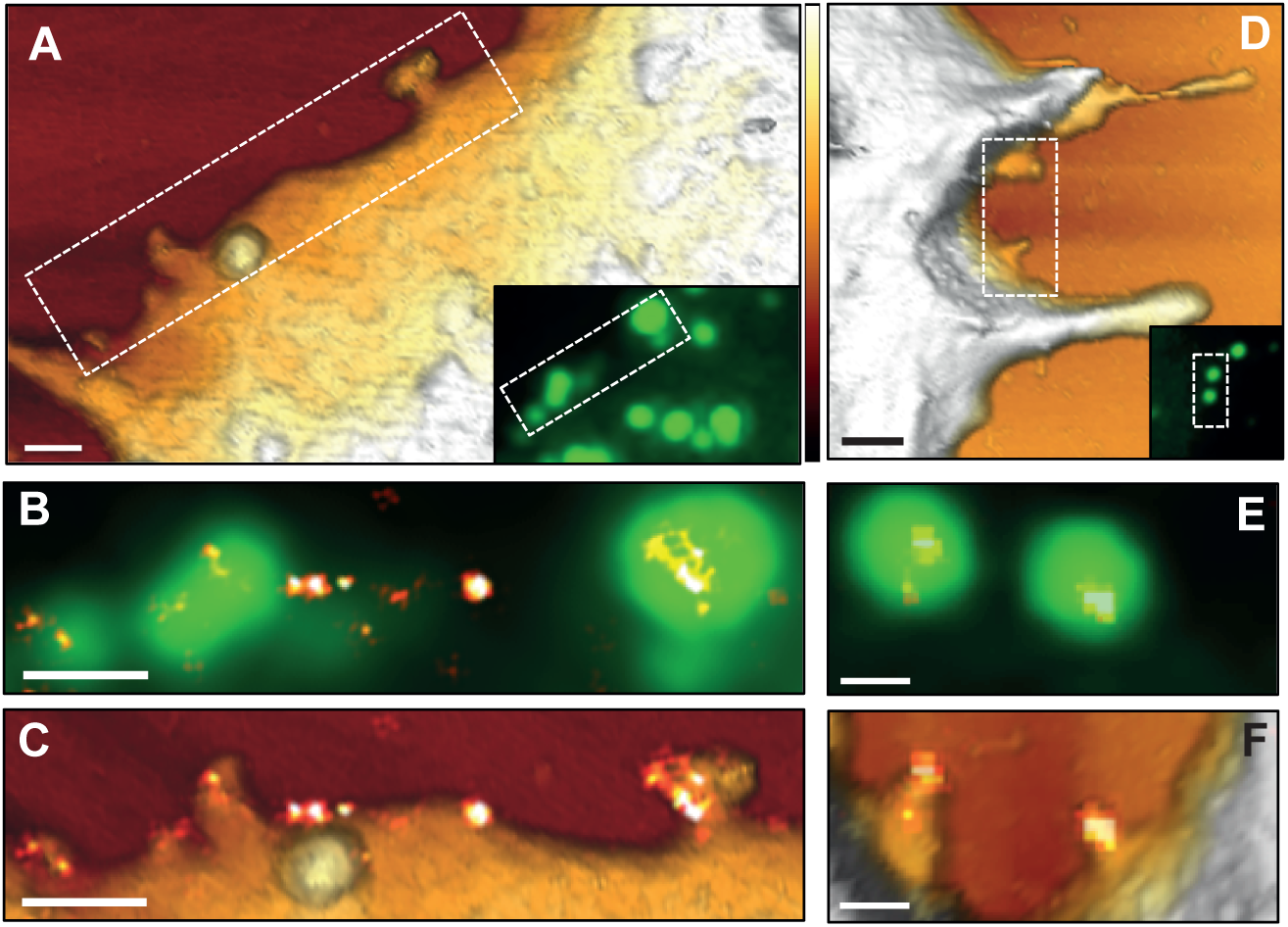
CD9 recruitment at HIV-1 budding sites. HeLa cells expressing HIV-1 Gag-GFP were immuno-stained with anti-CD9 coupled to Alexa-647 and imaged by AFM (first row), conventional fluorescence (second row) and dSTORM (third row) TIRF illumination: A and D) AFM 3D images of two different cells (the dotted line delineates the zoomed areas shown below and the insets are the corresponding Gag-GFP signal fluorescence images); B and E) overlays of the Gag-GFP picture with the reconstructed dSTORM image of the tetraspanin CD9. Scale bars are 500 nm (A, B, C and D) or 200 nm (C and F). The colour z scale shown in A and D is 300 nm.

**Figure 5.**
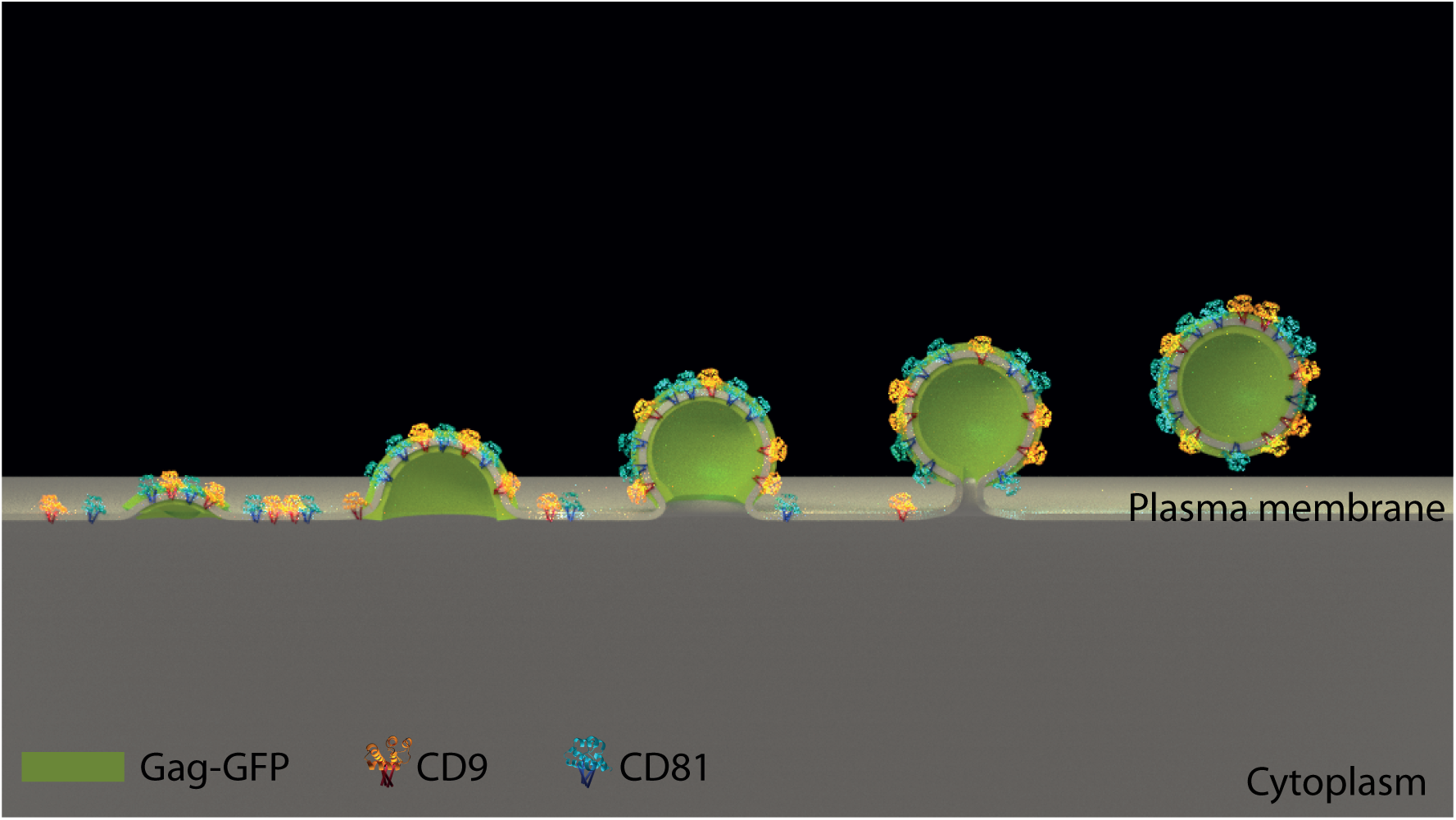
Model of tetraspanin lateral organization in the HIV-1 context. This scheme represents how the tetraspanins CD9 and CD81 (in blue and yellow) are laterally segregated within Gag-induced budding sites (in green). This leads to decreased CD9 and CD81 levels in the surrounding plasma membrane (in light grey), resulting in a net protein loss when Virus-Like Particles are excised.

In order to minimize the drawback coming from the calculation of CD9 densities of localization detected in 2D projected areas that could lead to an overestimation of CD9 densities in membrane buds, we calculated the actual CD9 density as the number of dSTORM localizations divided by the membrane area extracted from AFM images (see supplemental Materials). Taking into account the membrane topography, we confirmed that CD9 density is higher in Gag-GFP budding sites (3836 ± 934 localizations/μm^2^) compared to membrane regions where Gag proteins are absent (505 ±111) (Fig. S2B).

## DISCUSSION

Despite considerable progress in understanding the molecular mechanisms of HIV-1 replication cycle, several key aspects have not been yet fully addressed. In this study we investigated how HIV-1 subvert plasma membrane components using a combination of AFM and dSTORM. We report how HIV-1 Gag affects the lateral organization of the CD9 tetraspanin that becomes trapped within the viral particle.

We characterized CD9 localization in control cells using dSTORM and reported a clustered organization. Other tetraspanins have been previously characterized by high-resolution techniques, namely STED (36) and dSTORM (37,38). Interestingly, their localizations were also clustered, yet with slightly larger sizes (100-150nm wide) than found here for CD9 (67 nm). However, even though CD9 transiently associates with the tetraspanins characterized in these studies (e.g. CD82, CD81, CD53), their distribution is not expected to be identical. In fact, the area of CD81 clusters measured in the present work was slightly larger than that of CD9 (disk of 81 nm diameter, Fig. S5). This difference fits well with the slower diffusion of CD81 compared to CD9, as measured by single molecule tracking (39). It is interesting to note that cluster sizes measured in the publications described above fall in the same range and support a model whereby tetraspanins diffuse in the plasma membrane, embedded in small assemblies that could contain other tetraspanins, some protein partners, and lipids (10,36).

CD9 and CD81 being recruited at budding sites during HIV-1 egress suggested that functional platforms could originate from the gathering of these small assemblies. Indeed we confirm here that, upon expression of Gag-GFP in a model system recapitulating HIV-1-induced VLP production, CD9 and CD81 lateral organization is dramatically changed (respectively Fig. 2 and Fig. S4B) and the two tetraspanins are highly concentrated in regions of Gag assembly, forming large tetraspanin-enriched areas. Surprisingly the average cluster size did not radically change upon Gag expression (Fig. 2C and Fig. S4B), suggesting that the large CD9 and CD81 assemblies observed in Gag-enriched areas are composed of tetraspanin clusters that gather but do not fully coalesce (they can be resolved by dSTORM). This local concentration of tetraspanins is compensated by a decrease in their density in surrounding regions (as compared to control cells, Fig. S1E), indicating that CD9 enrichment in Gag+ areas is due to lateral reorganization of CD9 rather than protein recruitment from intracellular compartments. This model is in good agreement with our previous work showing that CD9, which diffuses in a Brownian motion in the plasma membrane, is trapped within Gag assembly sites (21). Gag is most likely the driving force that concentrates tetraspanin clusters, which probably cosegregate with other membrane partners that could facilitate budding. As previously suggested, lipid composition of these membrane assemblies could also play a key role in building large membrane platforms since Gag proteins are recruited by PIP2 lipids in a cholesterol-dependent manner (4,40). Correlative dSTORM / AFM on cells expressing Gag-GFP showed that CD9 concentrates at the very tip of nascent VLPs with almost complete exclusion from buds necks at late stages. Tetraspanins are thus most likely not involved in bud scission. But CD9 seems to have the propensity to partition within positively curved membranes. Such a behavior was previously described for Influenza virus where CD81 was mainly localized at the tip of viruses during late budding stages using EM (41) and is in good agreement with the presence of CD9 and CD81 in tubular structures such as membrane protrusions and filipodia (data not shown and (42). The correlation between CD9 density and membrane curvature could either reflect a role of CD9 in membrane remodeling and deformation, especially positive curvature, or its sensitivity to membrane curvature. At that time it is difficult to discriminate between these two models but the first hypothesis is more likely since we have also observed CD9 and CD81 in flat membranes. It is then more tempting to speculate that these proteins are important in the formation of highly curved membranes encountered during virus budding, in the production of exosomes from multivesicular bodies (43), as well as in cell fusion process observed during the gamete fusion where association of CD9 with high membrane curvature regions has been reported previously in oocytes (44) or during myoblast fusion (45). The acute enrichment of CD9 and CD81 into Gag-induced budding sites questioned about the functional role of these proteins in membrane remodeling and/or bud fission during viral production and we then performed their single or dual depletion in HeLa and measured VLP production (Fig. S5). Similarly to what has already been described (24), CD9 and CD81 do not seem to be essential for VLPs release in HeLa cells. However, we observed that CD9 silencing led to an increase of CD81 expression within VLPs, and vice versa, suggesting that one tetraspanin could compensate for the loss of another. Interestingly, this effect was not observed in cell extracts, emphasizing a specific role of tetraspanins in viral particles. The other side of the coin of this redundancy is the difficulty to clearly establish their functional role.

Markus Thali's group has demonstrated that, despite their enrichment at viral exit sites, the overall levels of tetraspanins are decreased in HIV-1-infected cells. More specifically CD81 down-regulation in HIV-1 infected cells was explained by its degradation in proteasomal and lysosomal pathways. These processes were shown to depend upon HIV proteins Vpu and Nef (28). Here, we report that Gag-GFP expression could also trigger CD9 and CD81 depletion in the absence of Vpu and Nef. In addition, tetraspanin levels remain normal upon expression of a Gag mutant impaired in VLP release (31), suggesting that CD9 and CD81 depletion is directly linked to VLP release from host cells. Even if the fraction of cellular plasma membrane escaping through this process remains low (as assessed by unaffected levels of CD46), CD9 and CD81 global protein levels are impacted most probably because of their high concentration within Gag assembly sites. Interestingly, tetraspanin levels influence HIV- 1 life cycle at different stages and in opposite manners. Indeed, CD81 was shown to potentiate HIV-1 transcription (25,46) through its association to SAMHD1 (47), while overexpression of tetraspanins at the surface of virions, including CD9 and CD81, decrease their infectivity (24,25). Tetraspanins thus appear as key elements in the modulation of HIV-1 virulence.

Taken together our results shed new light on the involvement of CD9 and CD81 during HIV-1 egress using a new type of correlative microscopy that is suitable to investigate virus-host interactions, providing topographic details and molecular mapping at the nanoscale in native conditions. More generally such correlative microscopy appears as an outstanding technique to analyze membrane remodeling and partitioning that regulate cell life.

## ACKNOWLEGEMENTS

We acknowledge the support from France-BioImaging (FBI, ANR-10-INSB-04), the European Research Council Starting (ERC-Stg-260787), the Agence Nationale pour la Recherche (ANR-15- CE11–0023) and the GIS IBISA (Infrastructures en Biologie Santé et Agronomie). LF and DSP were recipients of the French Ministry of Education and Research. LF was a FRM fellow and SD salary was paid with a Sanofi-Pasteur contract. We are grateful to Zhanna Santybayeva for creating the cartoon in Figure 5, Markus Thali for providing the Gag-GFP plasmids, and Heiko Haschke (JPK company) for his technical help and helpful discussion.

